# *cis*-MoO_2_(BHAN)_2_ Complex: Its Role in the Protection of Radiation-Induced DNA Damage

**DOI:** 10.1101/2024.01.17.576028

**Authors:** Priyangana Deb

## Abstract

The synthesized molybdenum complex, [cis-MoO_2_(BHAN)_2_] (BHAN**=** *β*-hydroxy-*α*-naphthaldehyde), exhibits remarkable efficacy in safeguarding DNA against radiation-induced damage. Comparative studies reveal that the complex offers superior protection to radiolysed DNA compared to the ligand (BHAN). Notably, at a concentration of 2 mM, the complex demonstrates the capability to shield 90% of damaged plasmid DNA from a 20 Gy radiation exposure. Additionally, it also affords significant protection against radiation-induced damage to cellular DNA (CTDNA) from gamma rays. These findings underscore the significant potential of cis-MoO_2_(BHAN)_2_ as an effective radioprotector for normal tissues in the context of radiotherapy. The results of this study contribute valuable insights into the advancement of radioprotective strategies, presenting a noteworthy breakthrough with implications for future medical advancements.

## 1. Introduction

Radiation therapy is an essential treatment for many types of cancer, but it can also damage healthy tissue. This damage is mainly caused by reactive oxygen species (ROS) generated during radiolysis^1^, which can attack and modify DNA molecules. One potential strategy to mitigate this damage is to use antioxidants that scavenge ROS and prevent their harmful effects^2^. Dioxomolybdenum complexes have shown promise in this regard due to their ability to act as efficient radical scavengers^3^. These complexes consist of a molybdenum atom surrounded by two oxo-bonds in a *cis* configuration, with additional ligands that can influence their reactivity and selectivity. By interacting with ROS and converting them into less harmful species, dioxomolybdenum complexes could potentially protect DNA from radiation-induced damage. Recent studies have investigated the interactions between the Dioxomolybdenum complex and DNA and their ability to protect DNA from radiation damage^4^. In this chapter, the ability of the synthesized Dioxomolybdenum complex to protect DNA from gamma radiation is explored and demonstrated. It has been found that the complex was able to scavenge radicals generated during radiolysis which was examined by Electron paramagnetic rasonance (EPR) and thus can prevent DNA damage. The protective effects of Dioxomolybdenum complex has also been investigated *in vitro*. Overall, the Dioxomolybdenum complex shows promise as an efficient scavenger of ROS generated during radiation therapy. Thus, their ability to protect DNA from damage suggests that they could be used as radioprotectors for normal tissue during radiotherapy. Further studies are needed to optimize their selectivity and efficacy and investigate potential side effects or toxicity.

## 2. Experimental Section

### 2.1. Materials and physical methods

calf thymus DNA and supercoiled plasmid pUC19 DNA were obtained from Sigma Chemical Company, USA, and Genei, India, respectively. All DNA solutions were prepared in Tris-HCl buffer; at pH 7.4. Concentration of calf thymus DNA was determined considering molar extinction coefficient at 260 nm to be 6600 M^−1^ cm^−1^. Absorbance at 260 nm and 280 nm were noted for calculating A_260_/A_280_. Values being greater than 1.8 but less than 1.9 indicate the DNA was sufficiently free of protein. Quality of calf thymus DNA was also checked by its characteristic circular dichroism band at 260 nm recorded on a J815 Spectropolarimeter. The investigation was conducted using Millipore water.

### 2.2. Physical Measurements

Gamma radiation was passed through DNA solution with the help of a GC-900 Gama Chamber. Emission intensity were measured with a Perkin Elmer LS-55 spectroflourimeter. Electron paramagnetic resonance (EPR) measurements were performed in Jeol JES-FA 200 ESR spectrometer equipped with a Jeol microwave bridge. The gel electrophoresis study was carried out with UVP Bio Doc-It Imaging System and nicking was assayed by UVP DOC-ItLS software.

## 3. DNA binding Studies

### 3.1. Emission studies for DNA interaction

the Complex [*cis*-MoO_2_(BHAN)_2_] was also subjected to titration with increasing concentrations of DNA, by the emission spectroscopy technique. In this experiment, the excitation of the methanolic solution of *cis*-MoO_2_(BHAN)_2_ at 317 nm shows emission with λ_max_ at 353 nm. The experimental data obtained from fluorescence quenching were further analyzed using the Stern-Volmer equation (1)^3,5^, a widely used equation to quantify the quenching of fluorescence.

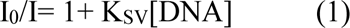

Where I_0_ and I are the fluorescence intensities in the absence and presence of DNA, respectively; and K_SV_ is the Stern–Volmer quenching constant, which is a measure of the efficiency of quenching by DNA, providing valuable insights into the nature and strength of the binding interaction between the *cis*-MoO_2_(BHAN)_2_ and calf thymus DNA. These results contribute significantly to our understanding of the binding behavior and potential applications of the *cis*-MoO_2_(BHAN)_2_ in DNA-related studies.

### 3.2. Viscometric study

Viscosity measurement data were presented as (η/η_o_)^1/3^ versus the ratio of the concentration of either of the ligand or the complex to that of the calf thymus DNA, where η_o_ is the viscosity of calf thymus DNA solution alone and η is the viscosities of calf thymus DNA solution in the presence of the *cis*-MoO_2_(BHAN)_2_^6^. The values were calculated from the observed flow time of calf thymus DNA by the relation; η = t−t_o_, where t and t_0_ are the values of flow times for the solution and the buffer respectively.

## 4. DNA damage and protection

### 4.1. Irradiation by gamma-ray

The calf thymus DNA and DNA-complex samples were incubated at 37 °C for 30 min and then irradiated in a ^60^Co γ-chamber at a dose rate of 1.88 kGyhr^−1^ for a total dose of 3.5 kGy. The irradiation doses for plasmid DNA samples were either 20 or 25 Gy. Our experimental design entailed administering a radiation dose that significantly surpassed the standard therapeutic dosage employed in normal radiotherapy to treat cancer patients. This deliberate augmentation of radiation dosage aimed to induce a comprehensive disruption of DNA, enabling us to evaluate the potential of the synthesized compounds in protecting DNA against such extensive damage. Notably, all experiments were meticulously conducted in well-controlled *in-vitro* settings, providing us with a suitable platform to explore the effects of higher radiation doses. This line of investigation holds promise for the clinical application of these compounds as good radioprotectors during radiotherapy.

### 4.2. Assessment of DNA damage followed by irradiation

#### 4.2.1. Estimation of radiation induced damage in calf thymus DNA by fluorescence spectroscopy

Here, calf thymus DNA solution was irradiated by a ^60^Co-γ source as described earlier. Then the radiation induced calf thymus DNA damage was assessed by fluorescence spectroscopy using ethidium bromide (EB) as probe. Thereafter, the emissions were recorded at 591 nm after excitation at 500 nm. This was compared with the fluorescence of the non-irradiated DNA-EB system under identical experimental setup. The dose–response relation is obtained from the plot of (I–I_a_)/(I_0_–I_a_) versus dose^3,7^, where I_a_ is the fluorescence intensity of EB, I_0_, the fluorescence intensity of EB-DNA control and I, the fluorescence intensity of EB-DNA irradiated sample.

#### 4.2.2. Protection of plasmid pUC19 DNA from radiation induced damage by the complexes and the ligand against different doses of gamma-radiation

Supercoiled pUC19 DNA solutions were pre-incubated for 30 min with different concentrations of complex or the ligand (0–2 mM) were exposed to different doses of 20 or 25 Gy gamma radiation. The agarose gel electrophoresis technique monitored the DNA damage protective ability of *cis*-MoO_2_(BHAN)_2_ and the ligand (BHAN). The samples were run on a 0.9% agarose in 1 X TAE buffer for 3 h at 80 mV, then it was treated with EB solution and the bands were visualized by UV light and photographed with UVP Bio Doc-It Imaging System.

#### 4.2.3. Mechanism of protection from radiation induced DNA damage by the ligands and the complexes; DPPH radical scavenging activity by EPR spectroscopy

Electron paramagnetic resonance (EPR) measurements were performed at room temperature (298 K). Spectroscopic parameters were 9.44 GHz (frequency), 100 mT (field sweep), 0.998 mW (microwave power) and a modulation amplitude of 3000 mT. The stability of freshly prepared DCM solution of DPPH was confirmed by monitoring^8,9^ the solution for 30 min, where no significant loss of signal was observed. In 100 µM DPPH solution, different concentrations (0–40 µM) of either *cis*-MoO_2_(BHAN)_2_ or the ligand (BHAN) were added and mixed thoroughly. The EPR signal was recorded after 2 minutes of mixing of either the complex or the ligand in appropriate cases with the DPPH solution under identical instrumental conditions.

## 5. Results & discussions

### 5.1. DNA binding studies

The emission intensity of the complex *cis*-MoO_2_(BHAN)_2_ exhibited a gradual decrease as calf thymus DNA was progressively added (Figure.1), which further supported the binding between the complex and DNA. The fluorescence intensity eventually reached a plateau at a relatively higher concentration of DNA, indicating that the binding reached saturation. This saturation point implies that the binding sites on DNA were fully occupied by the complex, resulting in a constant fluorescence intensity. To quantify the degree of fluorescence quenching, the Stern-Volmer equation was utilized, and the fluorescence quenching constant (K_SV_) was determined to be 1.6 x 10^4^ M^−1^ complex *cis*-MoO_2_(BHAN)_2_ at 25 °C. This value provides additional evidence for the binding of the complex to calf thymus DNA and highlights the efficacy of the complex in quenching the fluorescence emission. These compelling findings contribute to a comprehensive understanding of the complex’s interaction with DNA and its potential as a valuable tool in DNA-related applications.

**Figure 1:**
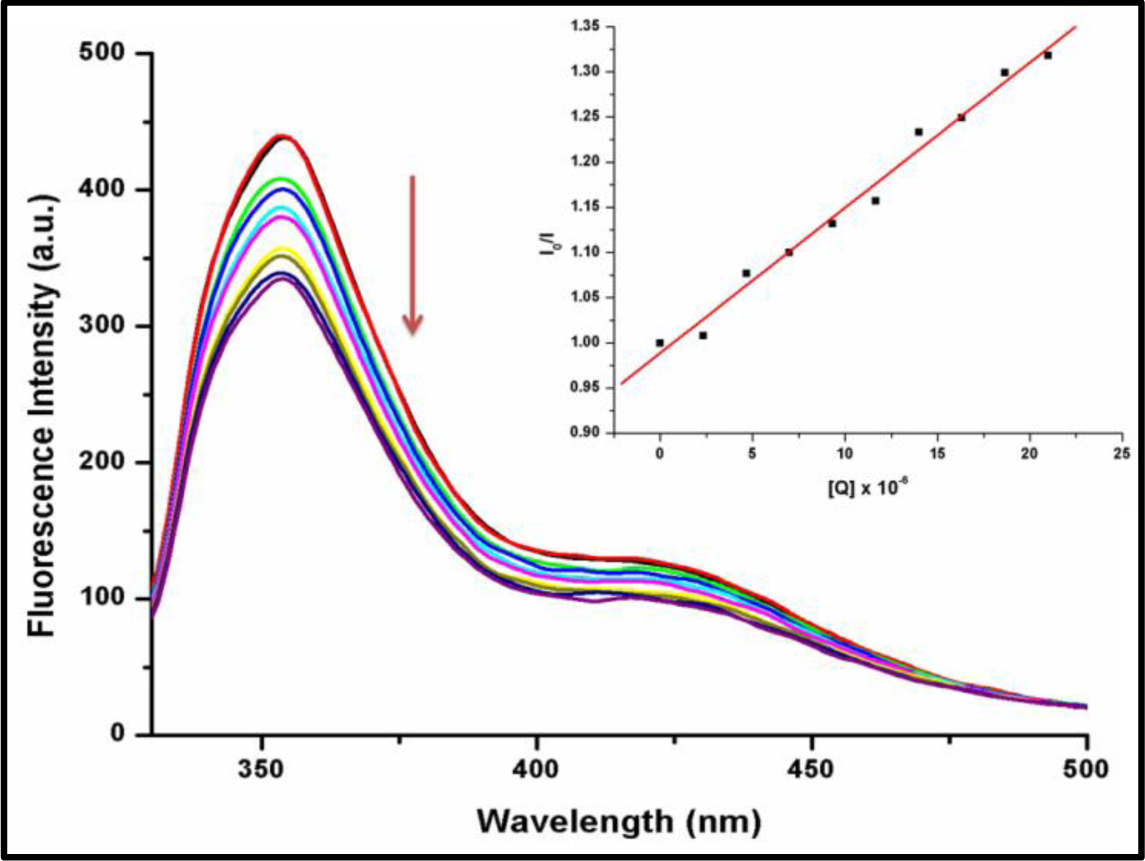
Emission spectra of ***cis*-MoO_2_(BHAN)_2_** in the presence of increasing amounts of DNA [Inset: Stern–Volmer plot]

Viscosity analysis, a highly sensitive technique, was then employed to gain insights into the mode of interaction between the complex and DNA. This method provides a deep understanding of the intricate nature of DNA binding^10–13^. The values of relative specific viscosities of DNA in the absence and presence of complex are plotted against [complex]/[DNA] (Figure.2). It is observed that the addition of *cis*-MoO_2_(BHAN)_2_ to the calf thymus DNA solution does not show any significant increase in the viscosity of calf thymus DNA, which clearly rules out the possibility of intercalation. If any small molecules bind to DNA through groove, that do not alter the relative viscosity value. Hence, in this case, the viscosity measurement results clearly hint at the groove binding of calf thymus DNA by *cis*-MoO_2_(BHAN)_2_.

**Figure 2:**
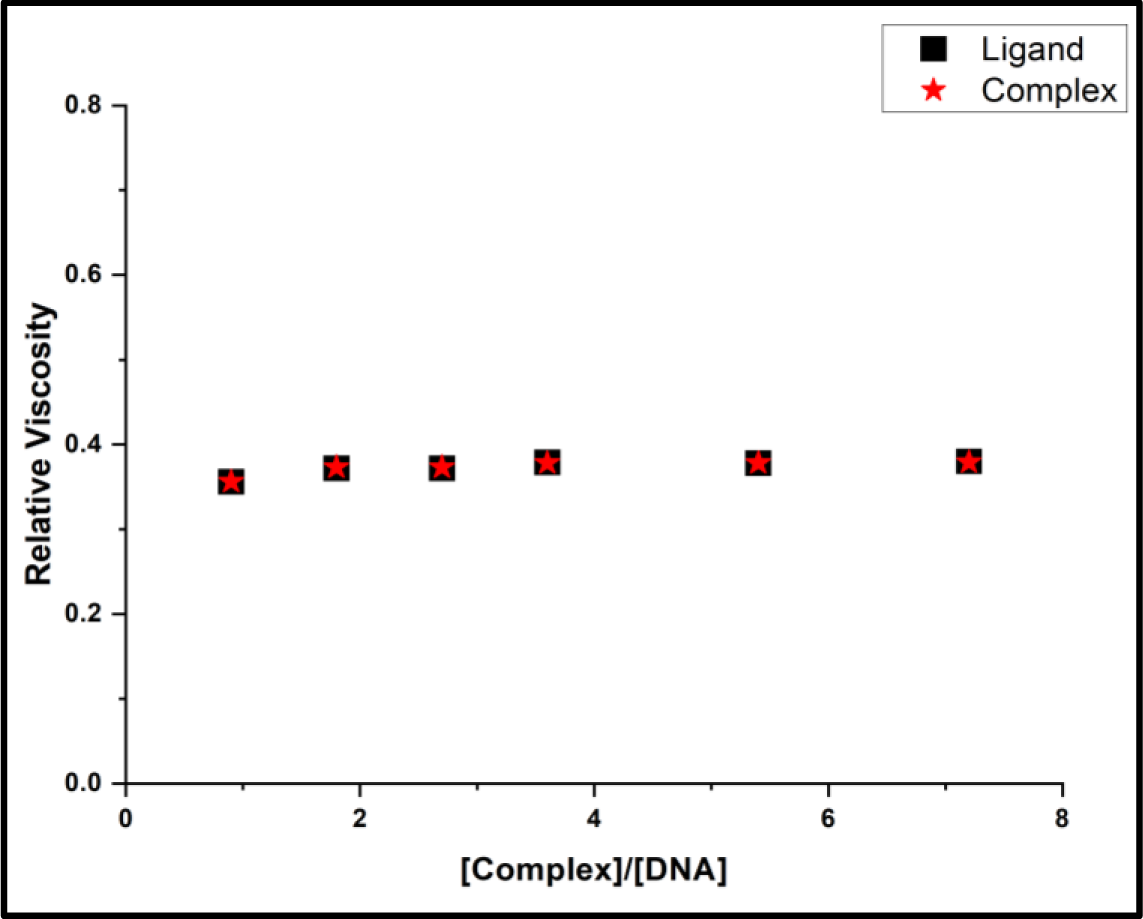
Effect of increasing amount of the **complex** [*cis*-MoO_2_(BHAN)_2_] and the **ligand** (BHAN) on the specific Viscosity of calf thymus DNA

### 5.2. Radiation induced DNA damage and protection

The assessment of DNA damage caused by radiation was conducted using a fluorometric technique that employed ethidium bromide (EB) as a probe^14–16^. EB has a weak emission in an aqueous solution, but when it binds with DNA, its emission intensity increases significantly due to its strong intercalation between adjacent DNA base pairs. If gamma irradiation causes damage to the DNA double helix then EB can’t intercalate with DNA, and this resulting damage of DNA due to irradiation, will cause relatively more EB to remain in the aqueous solution, which will be indicated by a reduction in fluorescence intensity. Therefore, this model was used to estimate radiation-induced DNA damage and evaluate methods for its protection^17^.

#### 5.2.1. Estimation of the protection of DNA damage by fluorometric technique

In order to evaluate the protective effects of a ligand or complex against radiation-induced DNA damage, DNA was pretreated with increasing concentrations of the ligand (BHAN) or *cis*-MoO_2_(BHAN)_2_ before exposure to radiation, after that EB was added to the solution. Then the resulting fluorescence intensity of EB-DNA solutions was measured, and the data is presented in Figure 3 (i)-(ii). The data showed that the fluorescence intensity of EB bound to irradiated DNA increased gradually with an increase in the concentration of the BHAN or *cis*-MoO_2_(BHAN)_2_, indicating that both the ligand and complex provide protection to the DNA against radiation-induced damage.

**Figure 3 (i):**
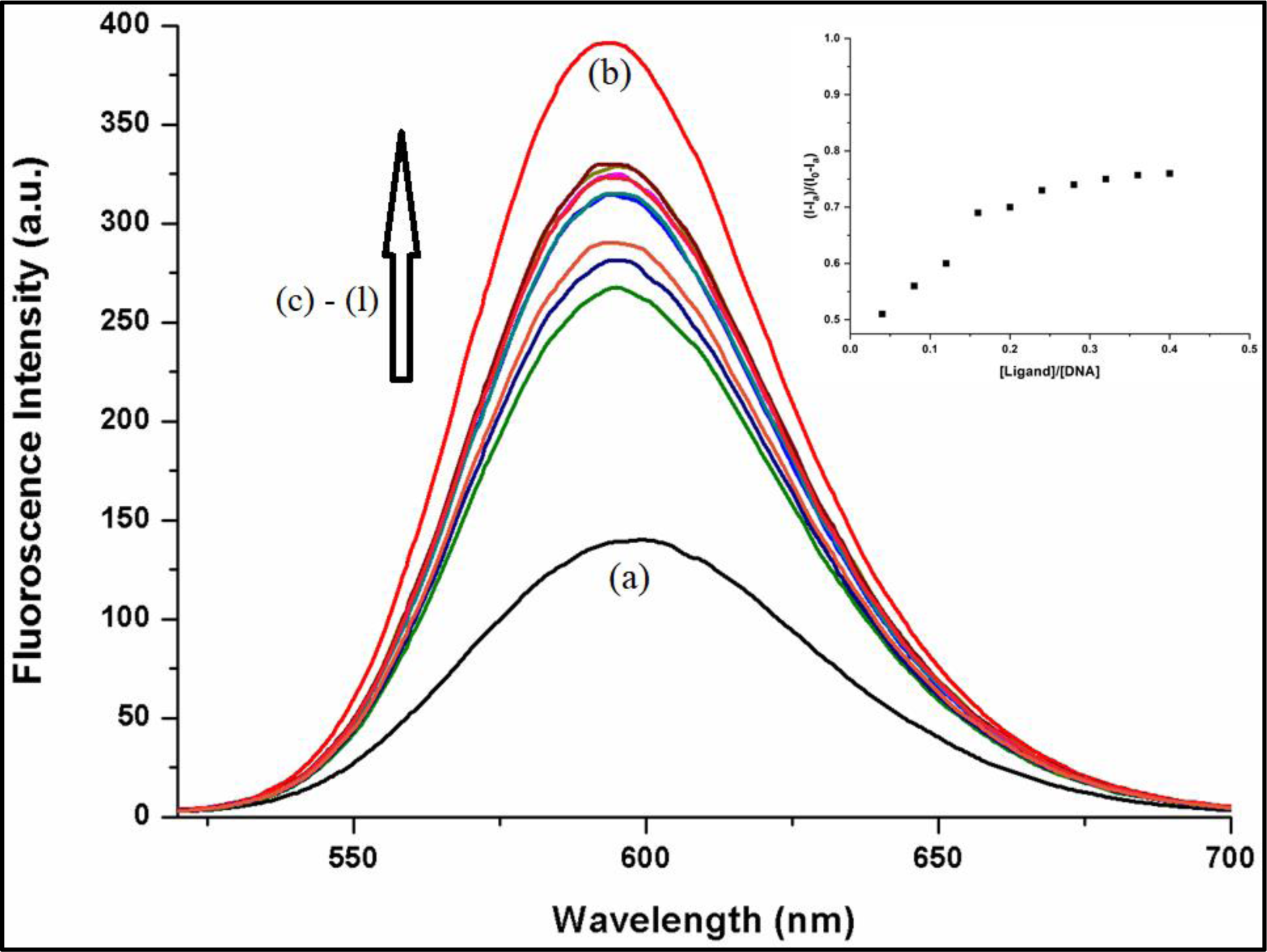
Fluorescence emission spectra of EB-DNA obtained after treatment of calf thymus DNA with EB following irradiation provided to DNA solutions either in absence or the presence of increasing amounts of **ligand (BHAN);** (a) = EB 900 μM, (b) = DNA 60 μM (unirradiated) + EB 900 μM (maintained 15 folds higher), (c)-(l) = DNA 60μM (irradiated) + increasing concentration of the ligand (BHAN) (0-24 μM) + EB 900 μM {Inset: Plot of (I − I_a_)/(I_0_− I_a_) versus [ligand]/[calf thymus DNA]}

**Figure 3 (ii):**
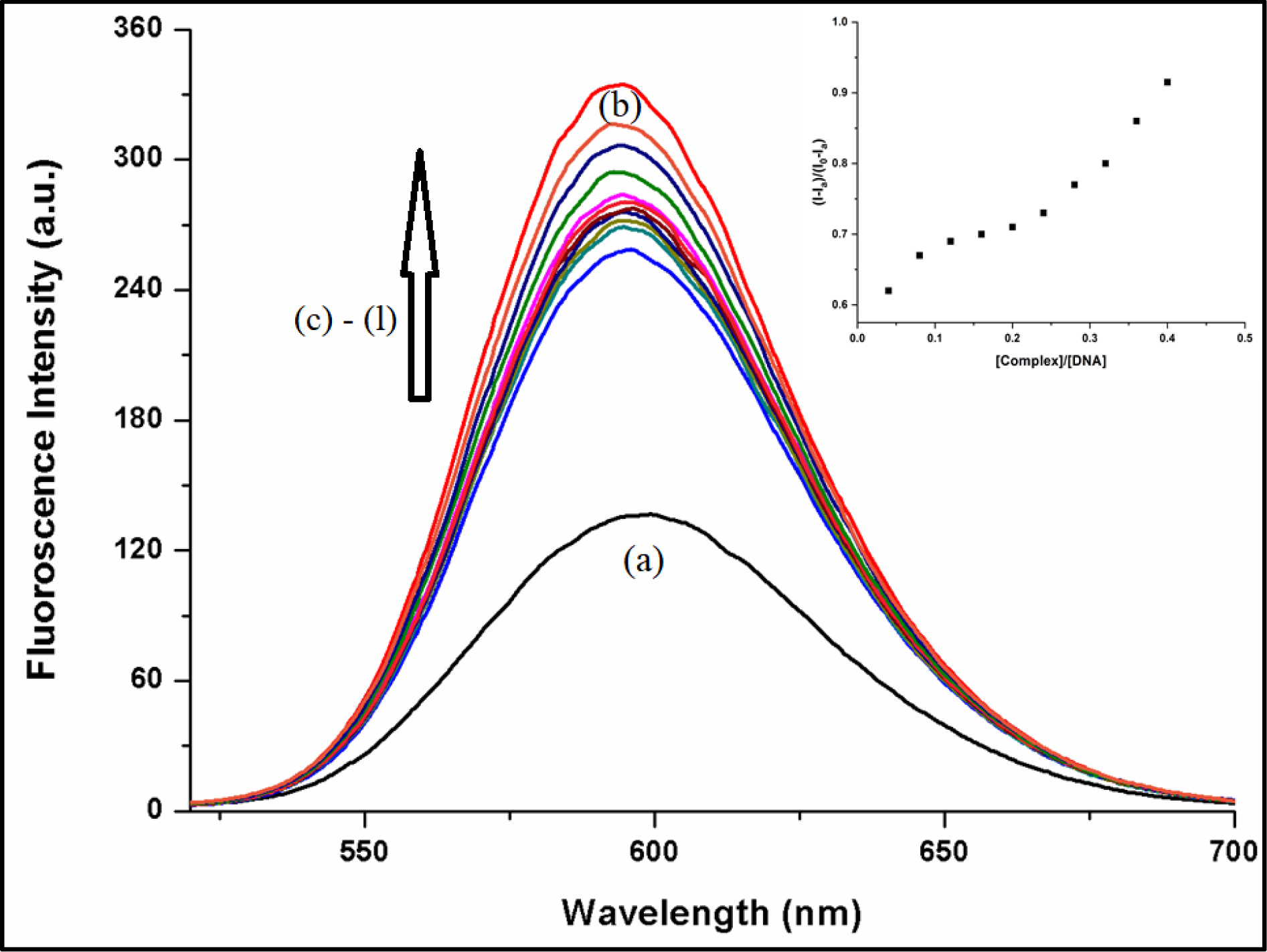
Fluorescence emission spectra of EB-DNA obtained after treatment of calf thymus DNA with EB following irradiation provided to DNA solutions either in absence or the presence of increasing amounts of ***cis*-MoO_2_(BHAN)_2_;** (a) = EB 900 μM, (b) = DNA 60 μM (unirradiated) + EB 900 μM (maintained 15 folds higher) (c)-(l) = DNA 60μM (irradiated) + increasing concentration of the complex (0-24 μM) + EB 900 μM {Inset: Plot of (I − I_a_)/(I_0_− I_a_) versus [complex]/[calf thymus DNA]}

Further a plot of (I − Ia)/(I_0_− Ia) versus [radioprotector]/[calf thymus DNA] was generated to calculate the percentage of protection exerted by either the complex or the ligand (shown in the inset of Figure 3 i & ii). The results indicated that as the concentration of the ligand (BHAN) or complex [*cis*-MoO_2_(BHAN)_2_] increased, the damage caused by radiation to DNA was inhibited. Specifically, the percentage of protection provided by the ligand (BHAN) was calculated to be 76%, while the complex [*cis*-MoO_2_(BHAN)_2_] can provide a much higher level of protection at 92%. The results indicate that the *cis*-MoO_2_(BHAN)_2_is highly effective in shielding the DNA from radiation induced damage. These findings have significant implications for the advancement of new approaches to minimize radiation-induced DNA damage in the future.

#### 5.2.2. The role of the ligand and complex in the protection from gamma-radiation induced strand breaks in plasmid pUC19 DNA

To further examine the protective capabilities of ligands and complexes against damage caused by gamma radiation on circular DNA, supercoiled pUC19 DNA was employed in a Tris-HCl/NaCl buffer (pH 7.2). The activity of the compounds was assessed using gel electrophoresis technique. Exposure of the plasmid DNA to gamma radiation at different doses (25 Gy and 20 Gy) resulted in strand breaks, leading to the relaxation of the plasmid DNA from a supercoiled (SC) form to a nicked coil (NC) form^3,14–16^, as seen in figures 4-5. However, when radiation-induced damaged pUC19 DNA was treated with varying concentrations of either the complex [*cis*-MoO_2_(BHAN)_2_] or the ligand (BHAN), significant protection from the damage was observed. The results of these experiments are presented in Table 1 (Figure.6 & 7).

**Figure 4:**
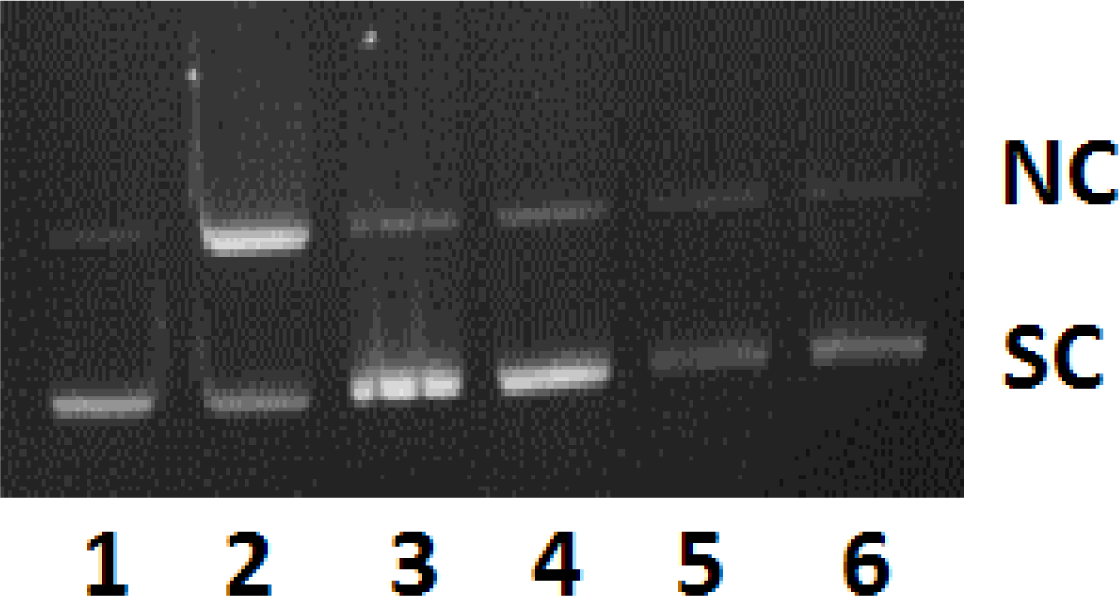
Protection of plasmid pUC19 DNA at **25 Gy** with different concentrations of **ligand (BHAN)** and ***cis*-MoO_2_(BHAN)_2_** on gamma-radiation induced strand breaks. *Lane1*: DNA control (No irradiation); *Lane 2*: DNA irradiated; *Lane 3*: DNA + 1 mM BHAN; *Lane 4*: DNA + 1 mM *cis*-MoO_2_(BHAN)_2_; *Lane 5*: DNA + 2 mM BHAN; *Lane 6*: DNA + 2 mM *cis*-MoO_2_(BHAN)_2_

**Figure 5:**
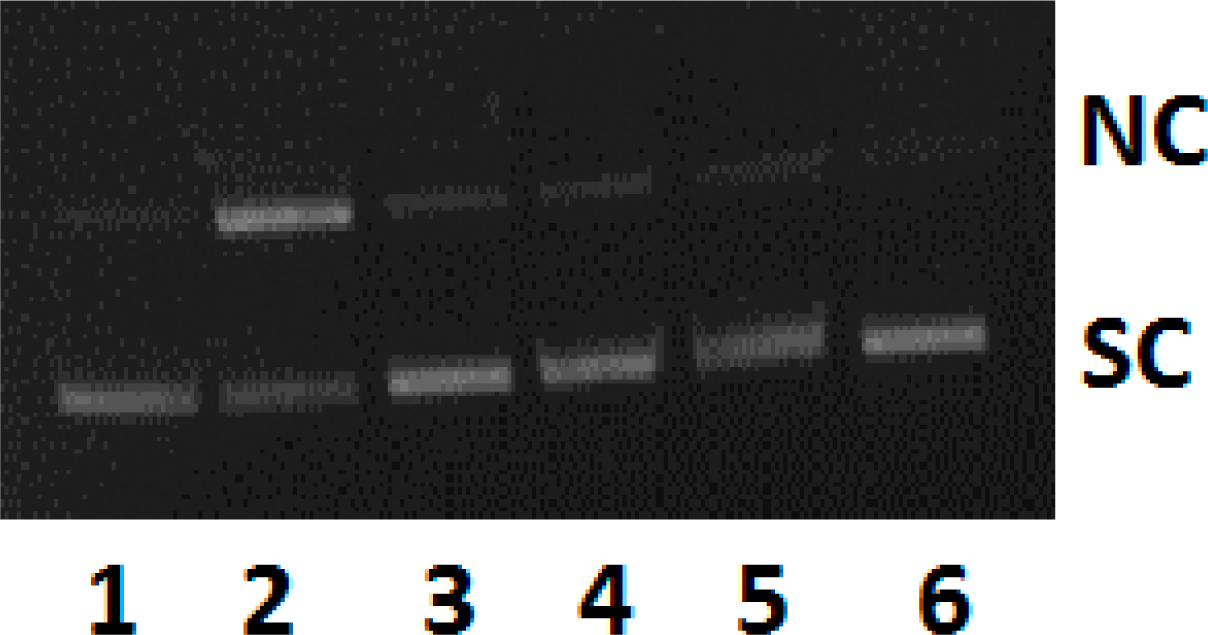
Protection of plasmid (pUC19) DNA at **20 Gy** with different concentrations of **ligand (BHAN)** and ***cis*-MoO_2_(BHAN)_2_** on gamma-radiation induced strand breaks. *Lane1*: DNA control (No irradiation); *Lane 2*: DNA irradiated; *Lane 3*: DNA + 1 mM BHAN; *Lane 4*: DNA + 1 mM *cis*-MoO_2_(BHAN)_2_; *Lane 5*: DNA + 2 mM BHAN; *Lane 6*: DNA + 2 mM *cis*-MoO_2_(BHAN)_2_

**Figure 6:**
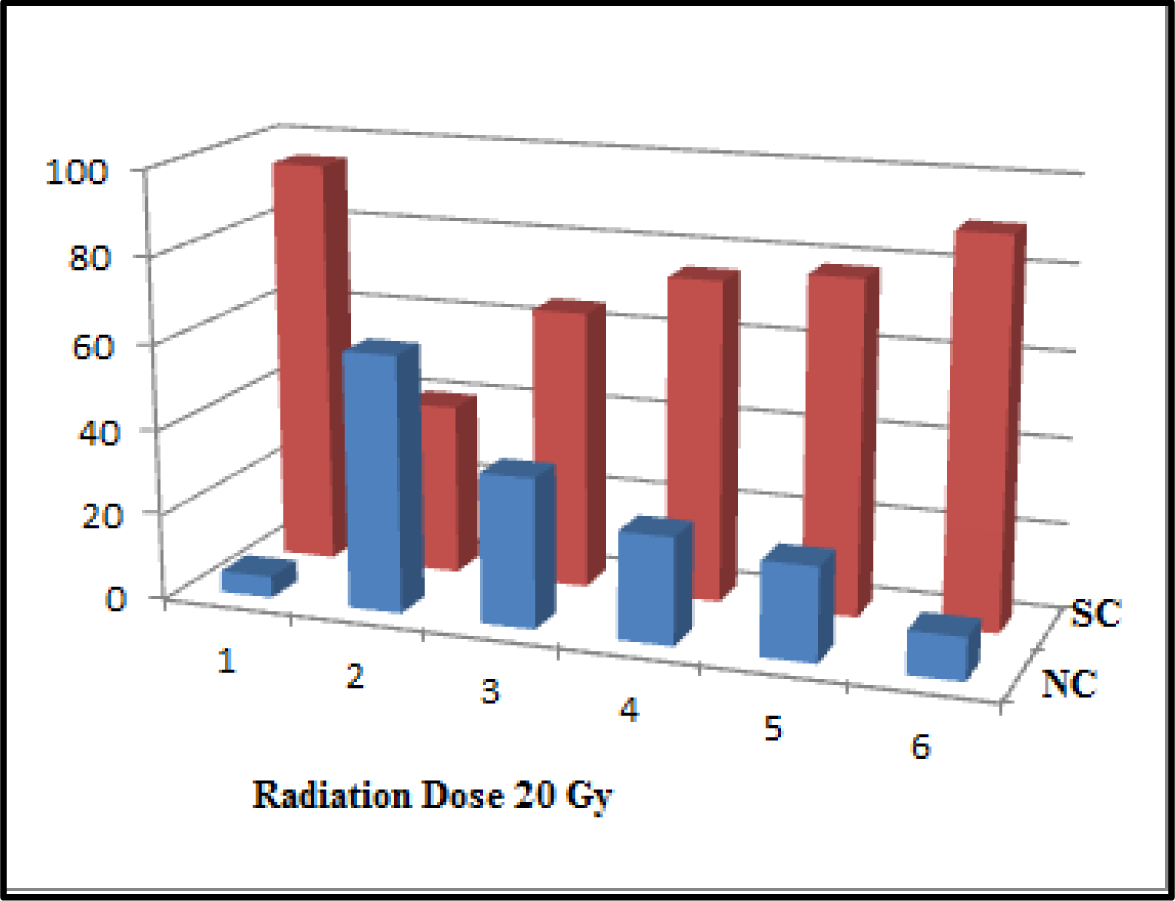
Graphical representation of protection by the **ligand (BHAN)** and the ***cis*-MoO_2_(BHAN)_2_** at radiation dose of **20 Gy**

**Figure 7:**
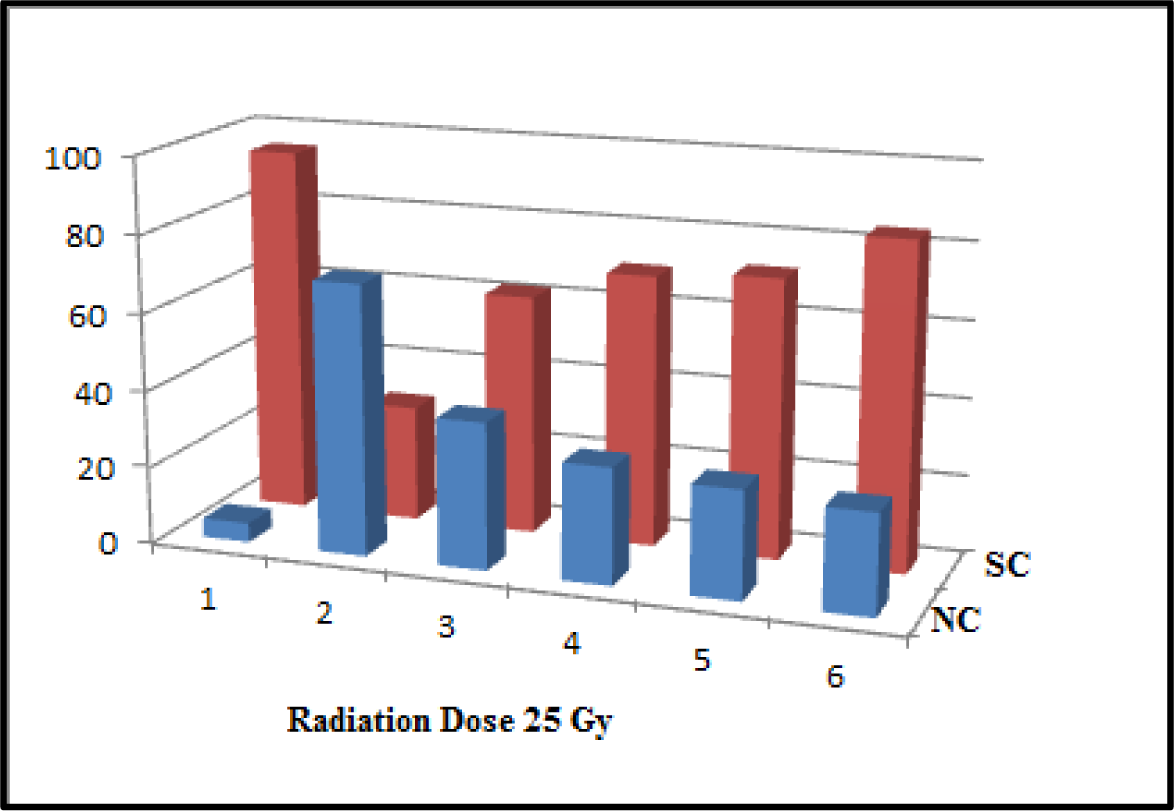
Graphical representation of protection by the **ligand (BHAN)** and the ***cis*-MoO_2_(BHAN)_2_** at radiation dose of **25 Gy**

**Table 1:**
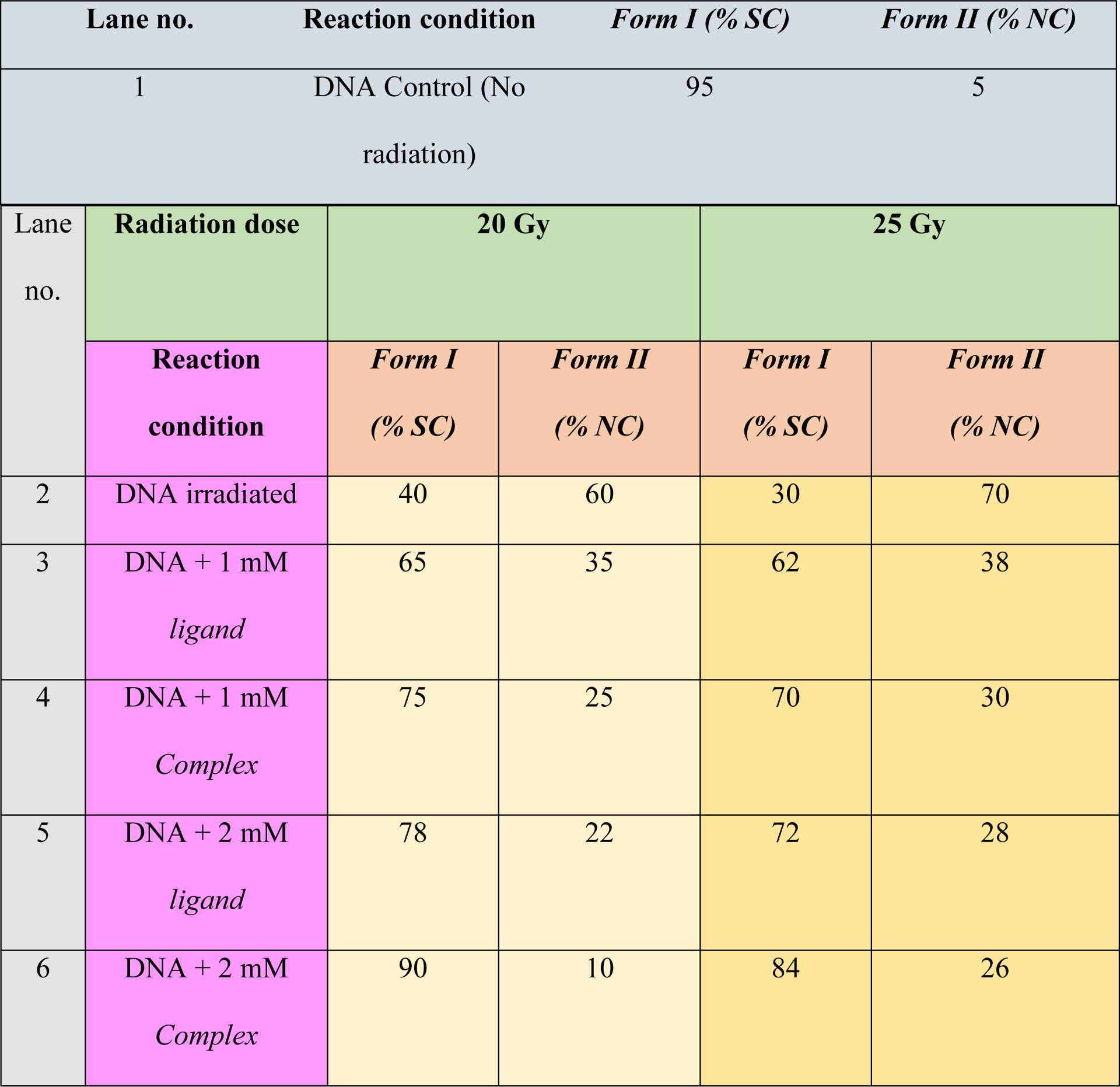
Extent of DNA SC pUC19 protection by the ligand (BHAN) and the *cis*-MoO_2_(BHAN)_2_.

The table provided displays the results of an experiment examining the effects of different treatments on the protection of DNA from radiation-induced damage. In Lane 1, the control group (without addition of any complex or ligand or any exposure to radiation) shows the percentages of Form I (supercoiled) and Form II (nicked circular) DNA, with 95% of the DNA remaining in the supercoiled form and only 5% in the nicked circular form.

In Lane 2 there is only plasmid DNA without addition of any complex or ligand and exposed to radiation doses of 20 Gy and 25 Gy. In this lane, the percentage of supercoiled DNA decreases while the percentage of nicked circular DNA increases which indicated the DNA damage due to radiation. Lane 2 shows 40% supercoiled DNA and 60% nicked circular DNA, with a radiation dose of 20 Gy, while with a radiation dose of 25 Gy, exhibits 30% supercoiled DNA and 70% nicked circular DNA.

To assess the protective effects of the ligand and complex, additional treatments were applied in Lanes 4 to 6. In Lane 4, the DNA was treated with a 1 mM concentration of the ligand, resulting in an increase in the percentage of supercoiled DNA to 75% and a decrease in nicked circular DNA to 25% only. Similarly, in Lane 5, the DNA was treated with a 1 mM concentration of the complex, resulting in 78% supercoiled DNA and only 22% of nicked circular DNA. Increasing the concentration to 2 mM for both the lgand and complex in Lanes 5 and 6 respectively, further enhanced the protective effects, with the complex demonstrating the highest level of protection. In Lane 6, the DNA treated with 2 mM complex exhibited 90% supercoiled DNA whereas nicked circular form is only 10%.

These results suggest that both the ligand (BHAN) and complex [*cis*-MoO_2_(BHAN)_2_] have a protective effect on DNA, with the complex demonstrating a greater ability to shield the DNA from radiation-induced damage. The findings highlight the potential of these compounds for mitigating radiation-induced DNA damage and provide valuable insights for future studies and the development of novel strategies in this field. The *cis*-MoO_2_(BHAN)_2_ can provide upto 90% and 84% protection against radiation doses of 20 and 25 Gy, respectively. The gel electrophoresis results clearly indicate that both the complex and ligand can offer significant protection to DNA against gamma radiation-induced damage in vitro by reducing the formation of NC form.

### 5.3. Mechanism of protection from radiation induced DNA damage; Assessment of the scavenging of DPPH by EPR spectroscopy

Radiation exposure in solutions can give rise to harmful radicals that pose a risk to biomolecules, including DNA. In order to mitigate this damage, it becomes crucial to eliminate the radicals generated during radiolysis. One effective strategy to protect molecules from radiation-induced harm involves the introduction of a compound with radical scavenging abilities into the solution during radiolysis. By doing so, this compound can effectively intercept and neutralize the harmful radicals. To evaluate the radical scavenging activity of both the ligand and the complex, electron paramagnetic resonance (EPR) spectroscopy was employed. EPR spectroscopy is a powerful technique widely used to investigate the behavior of radicals and their interactions with various compounds. In this study, the EPR signal of DPPH, a stable radical commonly utilized to assess the antioxidant capacities of natural and synthetic substances, was closely monitored^21^. This allowed for the accurate evaluation of the ligand and complex’s ability to scavenge radicals and protect against radiation-induced damage. The EPR spectra of DPPH in solutions of dichloromethane (DCM) were carefully observed during the titration process with incremental amounts of either the ligand (BHAN) or the *cis*-MoO_2_(BHAN)_2_. Remarkably, the intensity decrease observed in the spectra indicates that both the complex and the ligand possess the ability to effectively scavenge free radicals of DPPH (Figure.8). However, the impact appears to be more pronounced in the case of the complex. These compelling findings further reinforce the outcomes obtained from the experiments conducted to assess the protection against DNA damage.

**Figure 8(i):**
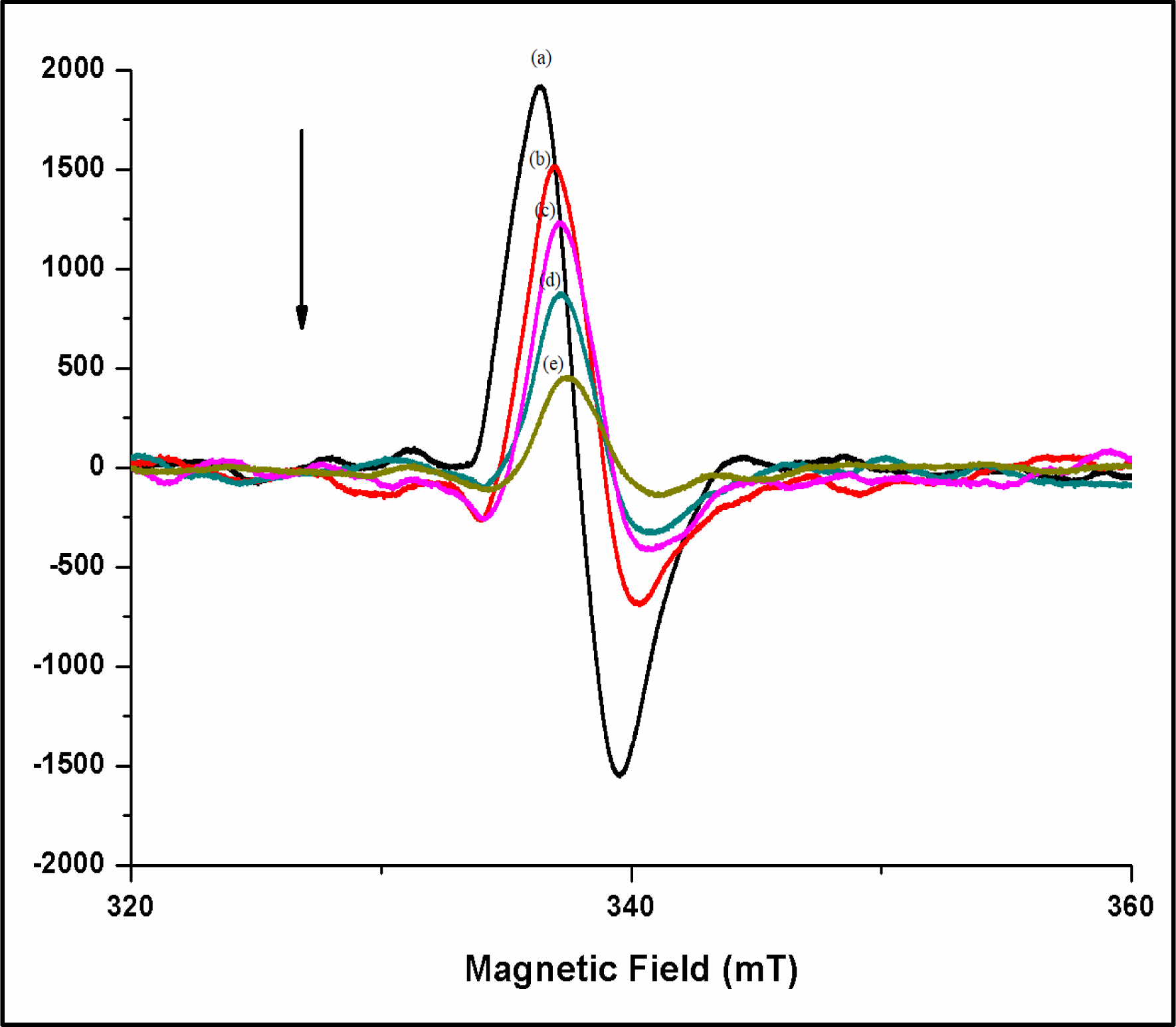
EPR spectra of DPPH (***a***:100 µM DPPH) with different concentrations of ***cis*-MoO_2_(BHAN)_2_** in DCM solution (**b**:10µM; ***c***:20µM; ***d***: 30µM; ***e***: 40µM)

**Figure 8 (ii):**
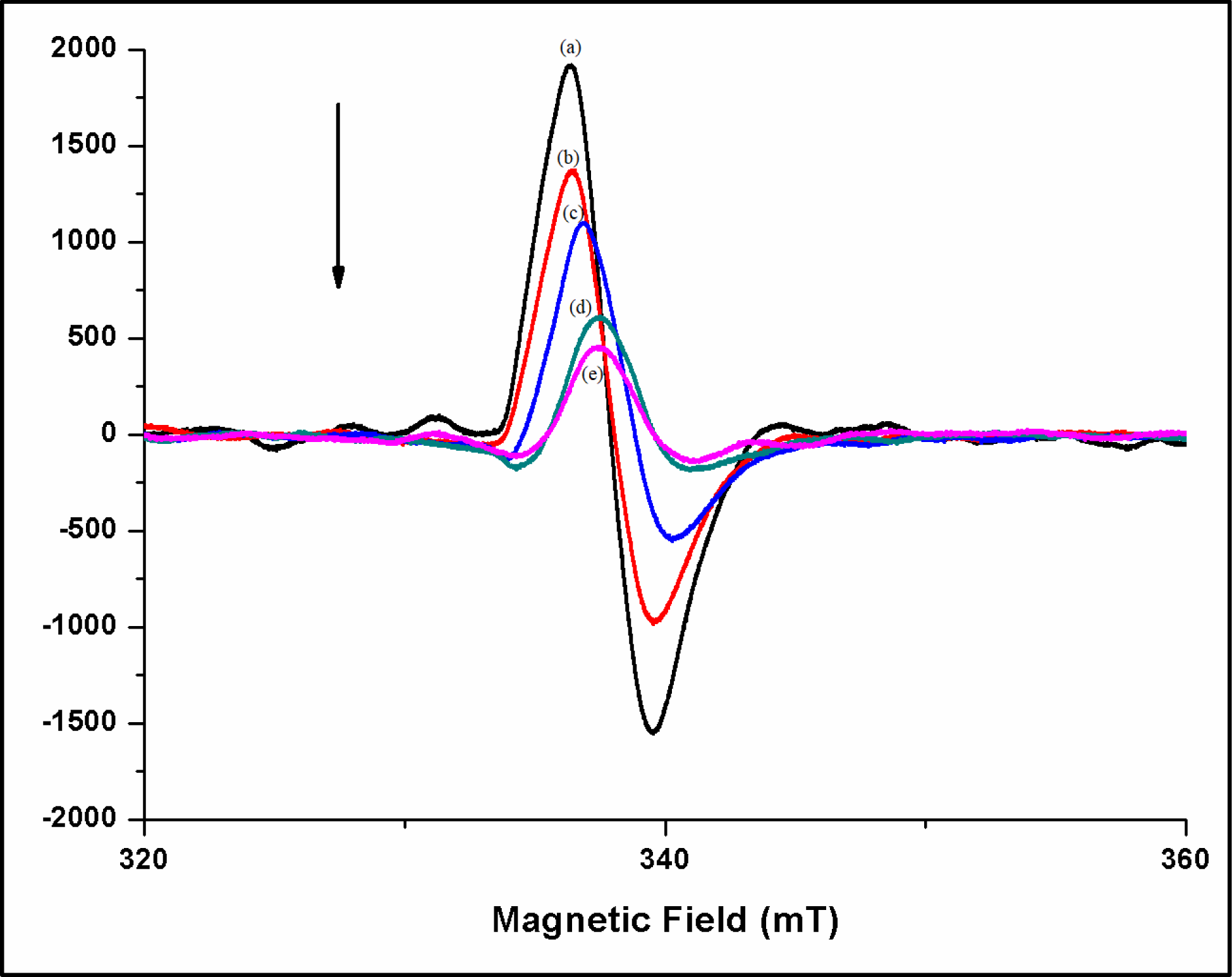
EPR spectra of DPPH (***a***:100 µM DPPH) with different concentrations of **ligand (BHAN)** in DCM solution (**b**:10µM; ***c***:20µM; ***d***: 30µM; ***e***: 40µM)

## 6. Conclusion

The molybdenum complex [*cis*-MoO_2_(BHAN)_2_] synthesized in this project has demonstrated a remarkable ability to protect DNA from radiation-induced damage. It was found that the *cis*-MoO_2_(BHAN)_2_ provided a higher degree of protection to radiolysed DNA than the ligand (BHAN). In fact, at a concentration of 2 mM, the complex was able to protect 90% of damaged plasmid DNA from radiation at exposure of radiation dosage of 20Gy. Furthermore, the complex was able to protect approximately 92% of radiation-induced damage to CTDNA from gamma rays. These findings suggest that the *cis*-MoO_2_(BHAN)_2_ has tremendous potential as an efficient radioprotector for normal tissues in radiotherapy. The ability of this complex to protect DNA from radiation-induced damage is particularly noteworthy, and the results of this study could have significant implications for the development of new therapies for a range of medical conditions. Overall, the complex’s ability to protect DNA from radiation-induced damage is a promising development in the field of radioprotection and could lead to important advances in the years to come.

